# Single cell RNA-sequencing of feline peripheral immune cells with V(D)J repertoire and cross species analysis of T lymphocytes

**DOI:** 10.1101/2024.05.21.595010

**Authors:** Raneesh Ramarapu, Judit M. Wulcan, Haiyang Chang, Peter F. Moore, William Vernau, Stefan M. Keller

**Affiliations:** Department of Surgical and Radiological Sciences, School of Veterinary Medicine, University of California Davis, Davis, CA, USA; Department of Anatomy, Physiology and Cell Biology, School of Veterinary Medicine, University of California Davis, Davis, CA, USA; Department of Pathology, Microbiology and Immunology, School of Veterinary Medicine, University of California, Davis, CA, United States; Department of Mathematics and Statistics, University of Guelph, Guelph, ON, Canada

**Keywords:** feline, T cells, single cell RNA-sequencing (scRNA-seq), T-cell receptor repertoire, cross species analysis, myeloid cells, V(D)J

## Abstract

**Introduction:** The domestic cat (Felis catus) is a valued companion animal and a model for virally induced cancers and immunodeficiencies. However, species-specific limitations such as a scarcity of immune cell markers constrain our ability to resolve immune cell subsets at sufficient detail. The goal of this study was to characterize circulating feline T cells and other leukocytes based on their transcriptomic landscape and T-cell receptor repertoire using single cell RNA-sequencing.

**Methods:** Peripheral blood from 4 healthy cats was enriched for T cells by flow cytometry cell sorting using a mouse anti-feline CD5 monoclonal antibody. Libraries for whole transcriptome, alpha/beta T cell receptor transcripts and gamma/delta T cell receptor transcripts were constructed using the 10x Genomics Chromium Next GEM Single Cell 5’ reagent kit and the Chromium Single Cell V(D)J Enrichment Kit with custom reverse primers for the feline orthologs.

**Results:** Unsupervised clustering of whole transcriptome data revealed 7 major cell populations - T cells, neutrophils, monocytic cells, B cells, plasmacytoid dendritic cells, mast cells and platelets. Sub cluster analysis of T cells resolved naive (CD4+ and CD8+), CD4+ effector T cells, CD8+ cytotoxic T cells and gamma/delta T cells. Cross species analysis revealed a high conservation of T cell subsets along an effector gradient with equitable representation of veterinary species (horse, dog, pig) and humans with the cat. Our V(D)J repertoire analysis demonstrated a skewed T-cell receptor alpha gene usage and a restricted T-cell receptor gamma junctional length in CD8+ cytotoxic T cells compared to other alpha/beta T cell subsets. Among myeloid cells, we resolved three clusters of classical monocytes with polarization into pro- and anti-inflammatory phenotypes in addition to a cluster of conventional dendritic cells. Lastly, our neutrophil sub clustering revealed a larger mature neutrophil cluster and a smaller exhausted/activated cluster.

**Discussion:** Our study is the first to characterize subsets of circulating T cells utilizing an integrative approach of single cell RNA-sequencing, V(D)J repertoire analysis and cross species analysis. In addition, we characterize the transcriptome of several myeloid cell subsets and demonstrate immune cell relatedness across different species.

## 1 Introduction

The domestic cat, Felis catus, is a valued companion animal and the second most popular pet in the United States. As of the American Veterinary Medicine Association (AVMA) pet ownership statistics, there are upwards of 46.5 million households with at least one cat in 2023 with ownership trends rising in the last decade from 25% in 2016 to 29% in 2022 (1,2). In addition to the growing need for feline veterinary care, the cat is an important model for infectious diseases such as virally-induced cancers (Feline leukemia virus, FeLV) and virally mediated immunodeficiency (Feline immunodeficiency virus, FIV) (3–6). Despite the value of the cat as a companion animal and infectious disease model, there are species specific research limitations including few cat-reactive immunophenotyping reagents and a poor knowledge base of immune cell markers and behavior.

In the advent of deciphering the feline immune system, many studies characterizing feline immune cells have relied strongly on antibody-based assays including flow cytometry and immunohistochemistry. This has allowed the study of immune cells in health and disease conditions such as FeLV or FIV (7–9). This approach depends on the assumption that feline immune cells are similar to other species. However, feline immune mediated diseases behave very differently from those in other small animals such as dogs, exhibiting unique etiologies and pathogenesis (10). In a cross-species context, innate immunity shows relatively high evolutionary conservation; however, this conservation dwindles with progression to adaptive immunity and complex immune phenotypes (11). Additionally, there is growing evidence demonstrating that evolutionary relationships do not translate to immune transcriptional or cell type relationships. Such an example is the mouse being a better model for human immune cells than the macaque (12). Thus, understanding the species-specific system in the context of our growing evolutionary database is desirable and there is a strong need to characterize the heterogeneous peripheral immune cell population in the cat, especially diverse populations such as T cells.

In efforts to investigate feline specific T cell diseases, our laboratory has contributed to the growing feline specific reagent and immune cell knowledge base including T cell receptor (TCR) repertoire analysis (13). In this trajectory, single cell RNA-sequencing is a logical next step. It enables the efficient large-scale capture and analysis of heterogeneous tissues, overcoming the challenges of species-specific reagent assays (14–17). Consequently, there has been a bloom in single cell RNA-sequencing studies of the peripheral leukocytes in a variety of mammalian species including the dog, cow, horse, and pig (18–22). Additionally, single cell data facilitates the study of cross species variations in immune cell type and evolution of these cells (12). ScRNA-seq of feline circulating leukocytes have been previously performed demonstrating the presence of 5 major cell types (T cells, B cells, NK cells, monocytes, dendritic cells) in the context of other non-model species (23). However, in-depth characterization of key subtypes and their marker conservation is yet to be determined. Thus, the goal of this study is to characterize the heterogeneity of circulating feline T cells and other captured leukocytes utilizing CD5 flow cytometry enriched scRNA-seq and V(D)J repertoire analysis in clinically healthy domestic shorthair cats between the ages of 6 months and 9 years. CD5 was chosen as a selective T-cell marker given the lack of availability of anti-feline CD3 antibodies. Additionally, we performed a cross species transcriptomic integration to assess feline T cells in the context of 4 other mammalian species - dog, horse, human, and pig. Our results provide a foundation for feline immune transcriptomics.

## 2 Results

### 2.1 Single cell atlas of the feline circulating immune cells

ScRNA-seq data obtained from 4 representative healthy cats of different ages consisted of 30,073 quality cells. These age groups were selected as they capture the major feline life stages: 6-month-old juvenile (6MO), 1-year-old mature (1YO), 4-year-old adult (4YO), and 9-year-old aged (9YO). The average number of cells captured per cat was 7,518 with an average of 3,655 transcripts per cell. **<Fig1A, Supplementary File1 S1A,B>**. Due to the poor annotation of the cat reference genome, we utilized a custom homologous mapping script based on ENSEMBL homologous mapping between the cat and human to improve gene readability. Our script increased the number of gene symbols mapped from 13,886 to 16,936, improving the quality of cell type assignment and biological interpretation. Following batch correction via reciprocal principal component analysis, unsupervised clustering revealed the presence of 21 clusters **<Fig 1B>**. A majority of cell types are equally represented from each sample **<Fig1C, 1D>**. A quality control violin plot for clusters can be found in the supplementary materials **<SUPPLEMENTARY File1 S2A>**.

**Figure 1.**
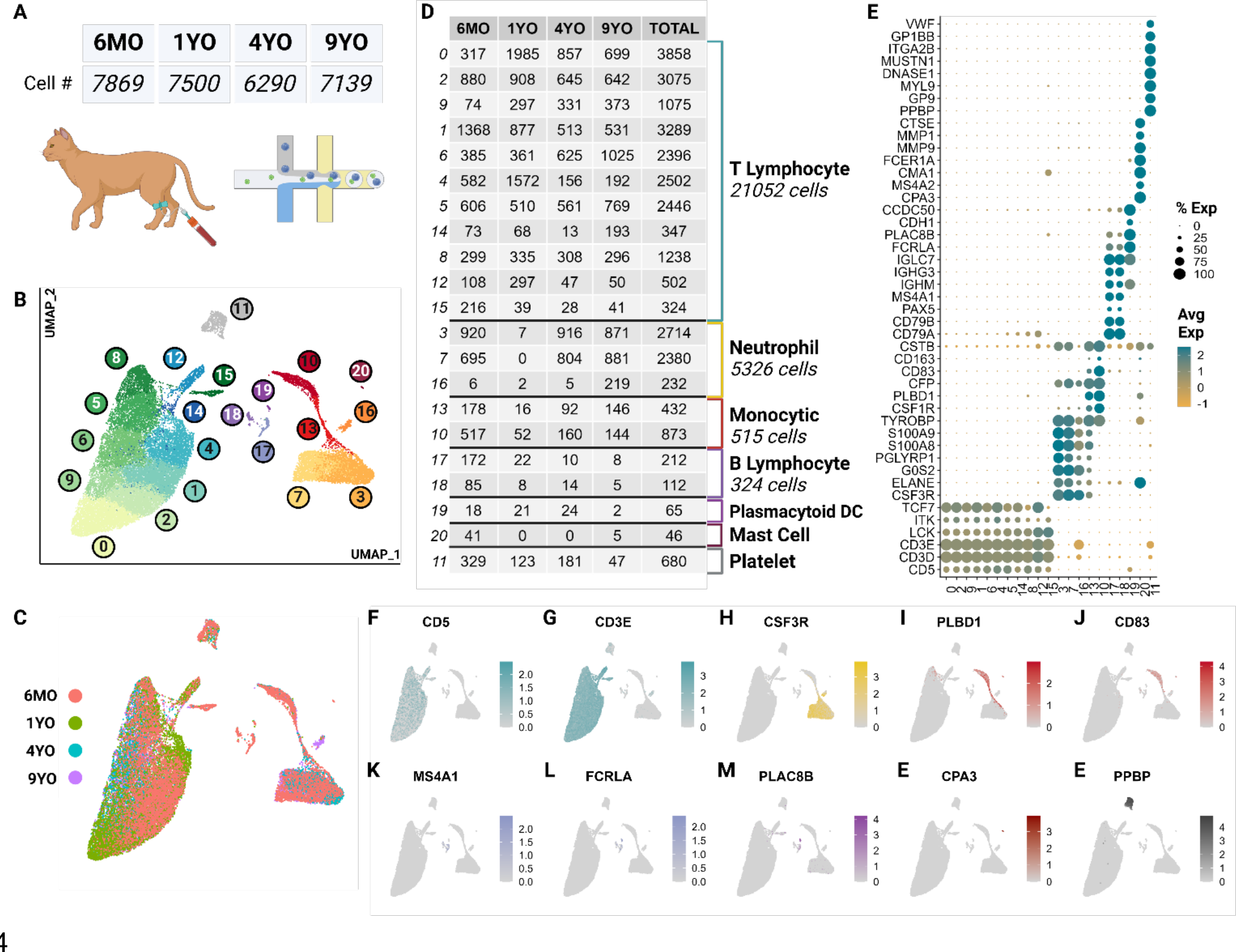
scRNA-seq atlas of CD5^+^ enriched circulating feline immune cells revealed 7 major types across 21 clusters. (A) Table of cell counts from each age group. (B) UMAP plot demonstrating the unsupervised clustering results for feline circulating immune cells from 4 healthy cats of different age groups. (C) UMAP plot of global clustering colored by age. (D) Table of cell type counts across clusters by age. (E) Dot plot demonstrating cell type specific marker expression of the 7 major cell types. (F-O) Feature UMAP plots demonstrating expression profile of key markers for each of the 7 different cell types.

These clusters were categorized into 7 major immune cell types based on canonical marker genes from multi-species PBMC single cell literature, top marker genes and over-enrichment gene ontology (GO) **<Supplementary File 2,3>**. The most numerous cell type identified was T cells called by high expression of T cell receptor complex genes (*CD3D, CD3E*) and species conserved T cell markers *LCK*, *ITK* and *TCF7* (12,23) **<Fig 1E-G, Supplementary File1 S2B>**. Across most T cell clusters, top identified GO terms of biological processes (BP) included T leukocyte processes of differentiation, migration, activation, and effector functions **<SUPPLEMENTARY File1 S3>**.

The second most abundant population was the neutrophils marked by neutrophil-derived proteases including elastase (*ELANE*), granulocyte colony stimulating factor receptor (*CSF3R*) and calprotectin (*S100A8/S100A9*) (24–26) **<Fig 1E,H>**. In most species, neutrophils are not captured in the PBMC layer during density centrifugation (27). Interestingly, our single cell data set captured a significant number of neutrophils from all but the 1YO sample **<Fig 1D>**. This is the first description of feline neutrophils at the single cell transcriptomic level and can provide insights into biological processes of these abundant cells.

A substantial number of monocytic cells were captured with marker genes including scavenger receptor for haptoglobin-hemoglobin complexes *CD163*, antigen presenting cell co-stimulatory molecule *CD83*, lipopolysaccharide detector protein *CD14*, phospholipase B domain containing 1 (*PLBD1*) and colony stimulating factor receptor (*CSF1R*) (28,29) **<Fig 1E,I,J>**. A relatively small but heterogeneous population of B cells were captured which were identified by conserved markers including B cell surface molecule *MS4A1* and Ig-alpha protein of the B-cell antigen component *CD79A* (23) **<Fig 1E,K,L>**. This population also expresses immunoglobulin genes (*IGHG3, IGHM*) indicating a sub-population of plasma cells (12) **<Fig 1E>**.

Unexpectedly, we captured plasmacytoid dendritic cells (pDC) and mast cells, which represented rare but discrete cell populations. pDCs were identified by expression of Fc fragment of gamma immunoglobulin (*FCRLA)*, dendritic marker *PLAC8B*, and immunoglobulin lambda constant 7 (*IGLC7*) and lack of expression of B cell canonical markers *MS4A1* and *CD79A* (30–32) **<Fig 1E,L,M>**. pDcs clustered more closely with B cells than other myeloid cells. This is due to close lineage associations and the shared expression of integral markers *FCRLA* and *IGLC7* among others (33) **<Fig 1E,L>**. Currently, there is debate regarding the reclassification of pDCs as innate lymphocytes and this analysis further supports such an argument (34) **<FIG1B>**. Mast cells were marked by IgE receptor *MS4A2* and a key component of mast cell protease, *CPA3* (35,36) **<Fig 1E,N>**.

Lastly, we captured a population of non-leukocytic immune cells which were annotated as platelets due to high expression of platelet specific proteins *PPBP*, *VWF* and *GP9* (37) **<Fig1E,O>**. This cluster also exhibited an extremely low average count of unique RNA and total RNA, which is consistent with a highly specialized cell type lacking a nucleus **<SUPPLEMENTARY File1 S2A>**.

### 2.2 Feline circulating T cells

To further characterize T cell subsets, we independently analyzed T cell clusters previously described at the global level (clusters 0,2,9,1,6,4,5,14,8,12, and 15). Unsupervised clustering of T cells revealed the presence of seven T cell clusters, which were unrecognizable at the global level **<Fig 2A>**. Largely, these clusters represent greater T cell phenotypes along increasing pseudotime values (proxy of differentiation or lineage) **<Fig 2B,C>**. The subtle transcriptional differences, as noted through overlapping differentially expressed genes, make effective annotation difficult **<SUPPLEMENTARY File1 S4>**. Nonetheless, we were able to identify several canonical subsets.

**Figure 2.**
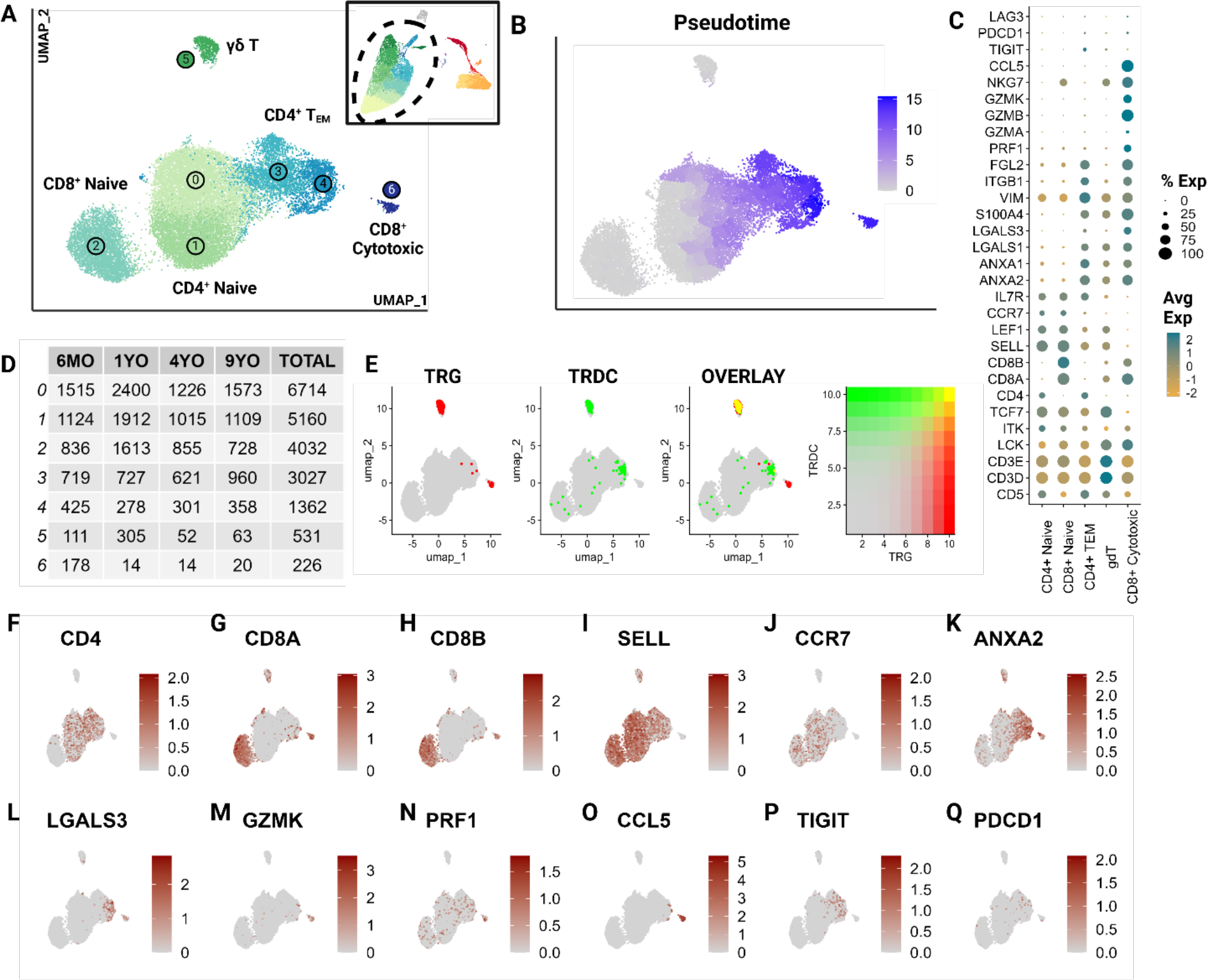
CD5^+^ enriched T cells segregate into naïve T cell subtypes and effectors. Unsupervised clustering of T cells reveals 7 subtypes. (A) UMAP of the scRNA-seq atlas of T cells. (B) UMAP of T cells colored by Pseudotime. (C) Dot plot of marker genes expressed by T cell type. (D) Table of cell type frequencies across ages by cluster. (E) Feature plots of expression and co- expression of *TRG* and *TRDC*. (F-Q) UMAP of T cells colored by classical T marker genes defining *CD4*/*CD8* status, naive (*SELL*, *CCR7*), effectorness (*ANXA2*, *LGALS3*, *GZMK, PRF1*), terminal differentiation (CCL5) and T exhaustion (*TIGIT*, *PDCD1*).

T cell subsets included *CD4^+^*, *CD8A^+^/CD8B^+^*and γδ T cells and were equally represented in all age groups **<FIG2D-H>**. A great majority were identified as naive cells, distinguished by their high expression of *CCR7* and L-selectin (*SELL*) (38–40) **<Fig2I,J>**. These naive populations segregated into two larger groups which could be distinguished by differential expression of *CD4* and *CD8A*/*CD8B* **<FIG2F-H>**. The *CD8A^+^/CD8B^+^* naive population represents a divergence from the traditionally observed distribution of naive T cells. Most naive T cells in mammalian species are *CD4*^+^ (12). However, a large population of *CD8A^+^/CD8B^+^*naive cells has been described in non-model species such as the horse (22). This highlights a potentially influential role of *CD8A^+^/CD8B^+^*T cells in cats.

The clusters 3 and 4, adjacent to the large naive populations, were identified as effector cells (TEM) **<FIG2A>**. These clusters show a gradual upregulation of T effector molecules including *S100A11,* galectins (*LGALS1, LGALS3),* annexins (*ANXA1, ANXA2*), and granzymes (*GZMA, GZMK*) (41,42) **<Fig2C,K-M>**. Szabo et al identified *CCL5* as a critical marker for highly differentiated effector T cells across tissues as well as its antagonistic expression to the naive marker *SELL* (41). In our data, Cluster 6 expressed the highest levels of cytotoxic markers including *CCL5* and exhibited the highest pseudotime score with no naive marker expression **<Fig2C,M-O>**. This indicates that this cluster is most consistent with the annotation of a terminally differentiated T cell cluster (CD8^+^ cytotoxic). Additionally, some effector cells expressed genes associated with T cell exhaustion such as *PDCD1* and *TIGIT* indicating the presence of a few of these cells in circulation <Fig2C,P,Q>.

Cluster 5 represented γδ T cells as inferred by the overlapping expression of the TCR genes *TRG* and *TRDC* (13) **<FIG2E>**. Within this population, there was an expression gradient of naive as well as effector markers, indicating subtypes within the γδ population **<Fig 2I,K,L>**. Although these T cell clusters represent biologically relevant groups statistically, the literature across various species identifies significant challenges associated with clustering of transcriptionally similar but heterogenous T cell populations (43). The γδ cluster exemplifies this challenge in that the cluster’s defining features overlap between two different but not mutually exclusive phenotypes - γδ recombination and effectorness/naiveness. These cells exhibit overlapping expressions of *TRG* and *TRDC*, indicating their TCR recombination-based categorization. An overwhelming majority of these γδ T cells are naive given their expression of naive marker *SELL* **<Fig 2I,J>**. However, some cells appear to express features of effectorness **<Fig 2K-N>**. This suggests some limitations of transcriptionally driven phenotyping of T cells. Nonetheless, this investigation represents a significant contribution towards deciphering the biology of feline T cells.

### 2.3 Feline circulating T effector cells

As described previously, subtyping T cells with overlapping phenotypes represents a significant challenge, which was further exacerbated by the low number of effector T cells in our dataset. Hence, we clustered effector T cells (clusters 3,4) independently, revealing 13 effector clusters **<FIG3A>**. Many of these clusters represent minimally divergent biological states as reflected by the marked overlap of top differentially expressed genes **<Fig3B, Supplementary File 5>**. However, direct visualization of known phenotypes demonstrate that the effector populations captured exhibit phenotypes of human TH1, TH2, TH17 and Treg cells **<FIG3C-J>** (41,42,44). Gene sets for each module calculation are provided in the supplementary materials **<Supplementary file 1, S3>**. These phenotypes do overlap and are not consistent with identified clusters. Nevertheless, the higher scores among closely positioned cells likely indicates the presence of these subtypes. However, a greater sample size of effector cells would be necessary to better resolve subtypes and their trajectory.

**Figure 3.**
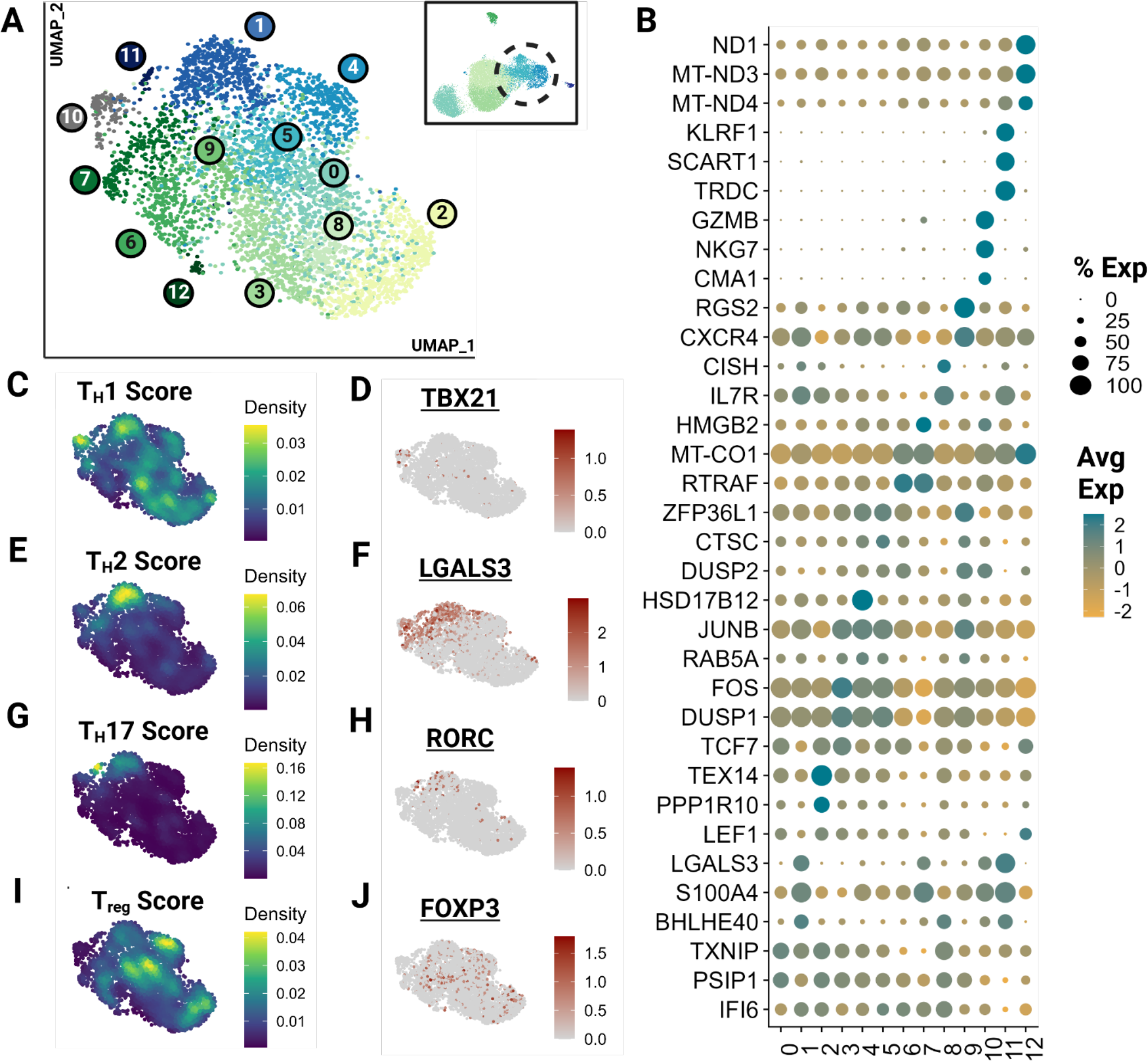
Feline Effector T cells (TEM) segregate into transcriptionally similar clusters and reveal the presence of helper T phenotypes. Unsupervised clustering of TEM reveals 13 subtypes. (A) UMAP of scRNA-seq atlas of TEM. (B) Dot plot demonstrating up to 3 top differentially expressed genes for each cluster determined via Wilcoxon rank sum testing (*Adj P <0.05*). (C,E,G,I) UMAP of TEM colored by helper T subtype gene modules. (D,F,H,J) Expression UMAP of a representative gene from each helper T gene module presented in parallel.

### 2.4 Cross species analysis of circulating T cells

To understand how the captured feline T cell populations compared to those of other domestic animal species and humans, we performed an integrative cross-species analysis. We selected 4 species for which peer-reviewed annotated scRNA-seq data with over 15,000 cells was available. ScRNA-seq data from three veterinary species (dog, horse and pig) and humans were chosen as the reference for homologous genome mapping (18,21,22,45). In total, we integrated 95,366 cells and identified 13 clusters across the 5 species **<FIG4A-F>**. These clusters were functionally annotated based on markers for T cell identity, naive phenotype, effector phenotype and cytotoxic phenotype in addition to their differential gene expression analysis **<Fig4G,H, Supplementary File 6>**.

**Figure 4.**
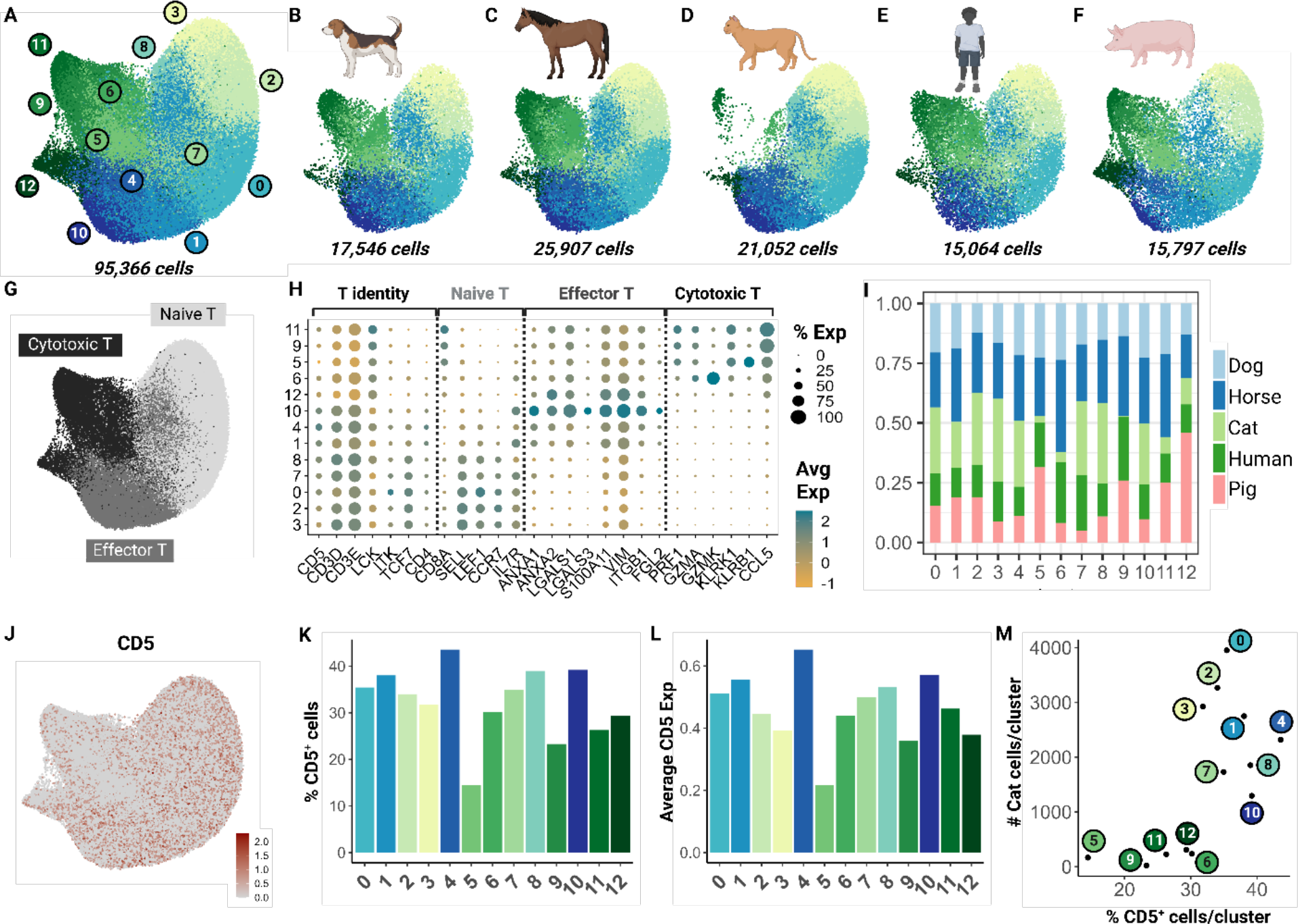
Cross-species integrative analysis of T cells reveals missing cytotoxic effectors in the cat. Unsupervised clustering revealed 12 clusters across 5 species. (A) UMAP of scRNA- seq atlas of T cells. (B-F) UMAP split by species- dog, horse, cat, human and pig. (G) UMAP of T cells colored by T cell phenotype. (H) Dot plot of marker gene sets for T cell subtypes. (I) Percentage bar chart of clusters stacked by species. (J) UMAP colored by *CD5* expression. (K) Bar chart of percentage *CD5^+^* cells per cluster. (L) Bar chart of average *CD5* expression across cells in each cluster. (M) Scatter plot of percentage *CD5^+^* cells per cluster versus number of cat cells within the corresponding cluster.

Although feline cells were overrepresented in some naive clusters such as 3 and 8, there was limited representation in clusters 5, 6, and 9 particularly **<FIG4I>**. To investigate whether this finding was associated with our enrichment protocol, we assessed *CD5* transcript expression across all clusters **<FIG4J>**. Our results suggest that the number of *CD5^+^*cells and average *CD5* expression were lowest in clusters 5 and 9 **<Fig4K,L>**. Additionally, we saw a direct relationship between the number of feline T cells and the *CD5^+^* cell count in each cluster. Particularly, clusters 5, 6, 9, 11, and 12 were most underrepresented **<FIG4M>**. These clusters were identified as cytotoxic effector cells **<FIG4H>**. This highlights a limitation of our CD5 enrichment protocol as it may select against specific effector clusters such as *CD8^+^* cytotoxic cells. Much of our current understanding of *CD8*^+^ feline T cells is from flow cytometric studies with no further subtyping due to lack of reagents. Thus, further investigation is necessary to validate whether this low frequency is biological or technical.

### 2.5 V(D)J recombination

To identify patterns in T cell receptor (TR) expression, we sequenced TR alpha (TRA), TR beta (TRB), TR gamma (TRG) and TR delta (TRD) transcripts simultaneously. The majority (>87.0%) of CD4^+^ naïve, CD4^+^ TEM and CD8^+^ naïve T cells expressed TRA/TRB transcripts only **<FIG5A>**. In contrast, only 35.0% of CD8^+^ cytotoxic T cells expressed TRA/TRB transcripts while 60.6% expressed TRA/TRB/TRG transcripts. γδ T cells primarily expressed TRG/TRD transcripts (49.7%) followed by TRG/TRD/TRB (16.4%) transcripts. Of note, 11.5% of cells had TRA/TRB/TRG transcripts, which might represent αβ T cells with an effector phenotype similar to gd T cells. CD8^+^ Cytotoxic T cells also exhibited differential TRAV gene usage, and to a lesser degree, TRBV gene usage **<Fig5B,C>**. The junctional length of TRG transcript in CD8^+^ Cytotoxic T cells was highly skewed towards 16 amino acids and converged on the conserved motif ‘CAAWDPRGYGWAHKVF’ **<FIG5D>**. Most TRG rearrangements involved cassette 2 (TRGV2- 2/TRGJ2-2) **<FIG5E>.** Interestingly, a subset of TRGV2-2 genes rearranged to the TRGJ2-1 gene, which is located upstream of TRGV2-2.

**Figure 5.**
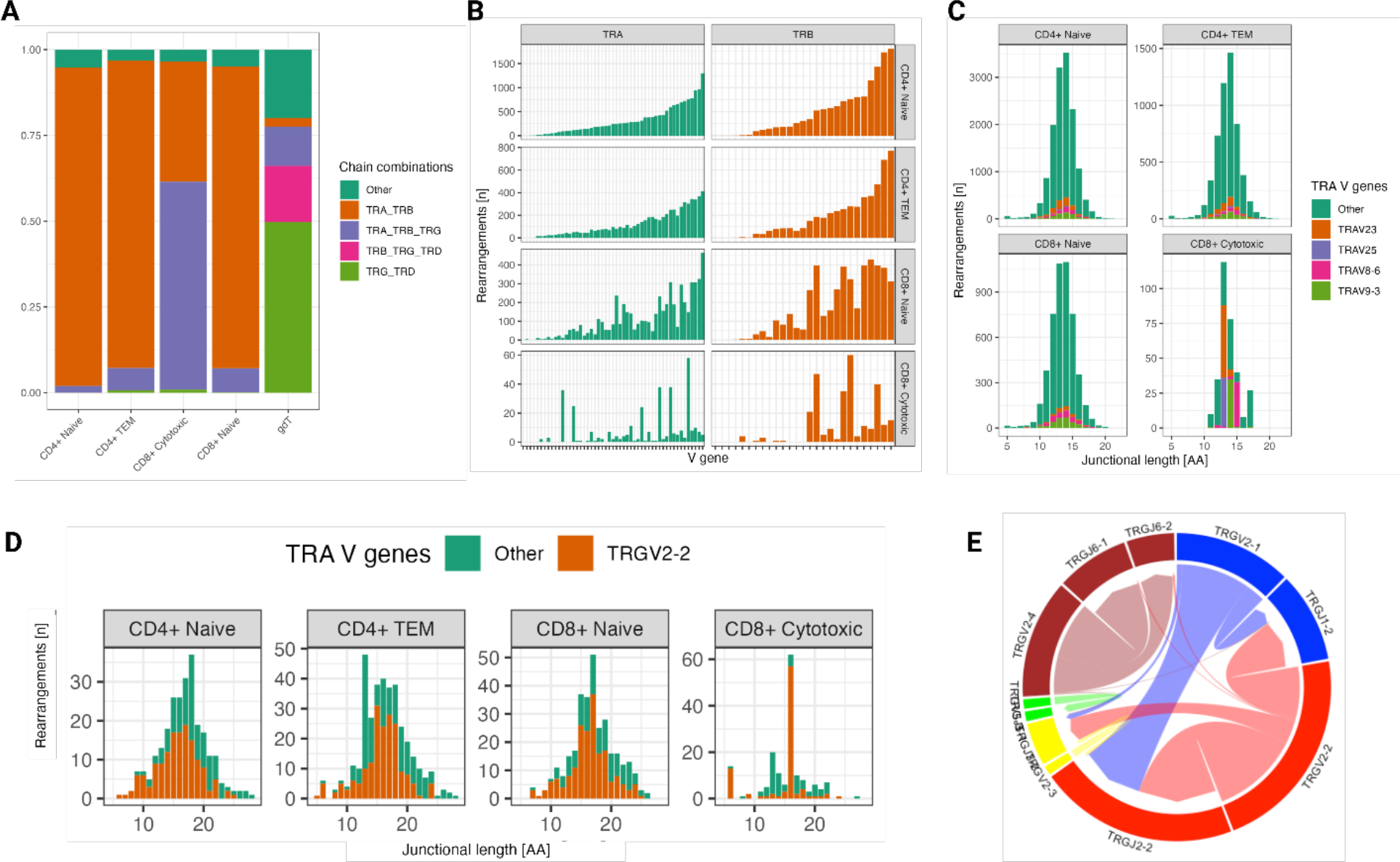
(A) Percentage of T cells expressing a certain combination of productive TR chain transcripts. TRA/TRB transcripts dominate in all ab T cell subsets except for CD8^+^ cytotoxic T cells, which primarily express TRA/TRB/TRG transcripts. (B) TRA & TRB V gene usage of the 4 major ab T cell subsets. CD8^+^ cytotoxic T cells show differential usage compared to the other T cell subset but are also substantiated by fewer cells. (C) Junctional length of TRA rearrangements of the 4 major ab T cell subsets. Junctional regions with dominant TRAV genes in CD8^+^ cytotoxic T cells are 13 and 14 amino acids long. (D) TRG transcripts in CD8^+^ cytotoxic T cells have a skewed junctional length with 16 amino acid length (top) and a conserved motif (bottom) (E) The majority of TRG rearrangements in CD8^+^ cytotoxic T cells involved cassette 2 (TRGV2-2/TRGJ2-2). Genes are displayed in order of genomic location and colored by cassette.

Next, we characterized the repertoire overlap between cats and the relatedness of junctional regions across cats and T cell subsets **<FIG6A>**. The number of shared clonotypes was highest for TRA and TRG but overall low **<FIG6A>**. When expanding the definition of shared repertoires to include clonotypes with one amino acid difference, the TRA locus had the highest number, size and publicity of clusters **<Fig6B,C>**.

**Figure 6.**
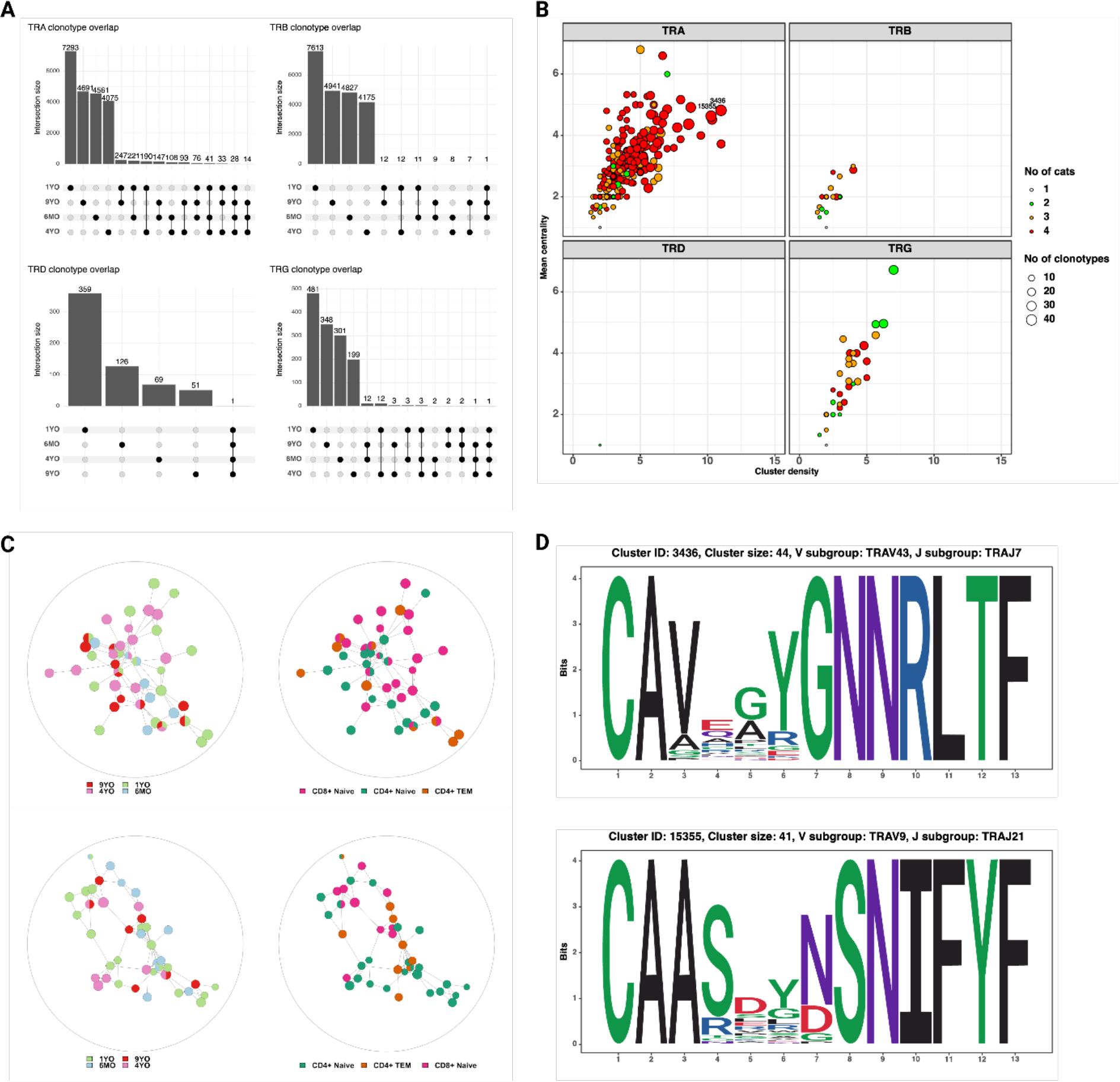
(A) Shared clonotypes across 4 cats. (B) Characterization of clusters based on mean centrality and cluster density. (C) Representative examples of TRA clusters with high centrality and density.

### 2.6 Feline circulating monocytic cells

Although CD5^+^ enrichment was performed for T cell selectivity, we serendipitously captured neutrophils and monocytic cells, which provides insight into the transcriptome of circulating myeloid cells in cats. Clustering analysis of both cell types was performed independently. Due to the low myeloid cell count in the 1YO, it was excluded from this analysis.

Monocytic cells clustered into 5 groups and were largely represented by the 9YO sample **<Fig 7A, Supplementary File 1, S4A>**. Cluster 3 represented neutrophils which is likely due to the close clustering of myeloid cells on the global object **<Fig 1B, Supplementary File 1, S4B>**. Cluster 0, 1 and 2 were identified as classical monocytes (CM) with all clusters having high expression of markers *PLBD1*, *CD83* and classical monocyte markers *CD14* and *VCAN* (28,29,46) **<Fig 7B-D>**. We did not identify any non-classical monocytes, based on absence of *CD16* expression, as seen in other species such as humans, horses and cattle (22,29,46,47). Cluster 4 showed an upregulation of receptor-type tyrosine-protein kinase *FLT3*, a conventional pan-dendritic cell marker (48) **<Fig 7F>**. In addition, these cells exhibited upregulation of MHC class-II molecules (*FECA-DRB*, *HLA-DRA*, *HLA-DOA*, and *HLA-DMD*) and were hence identified as conventional dendritic cells (cDC) (49) **<Fig 7G>**.

**Figure 7.**
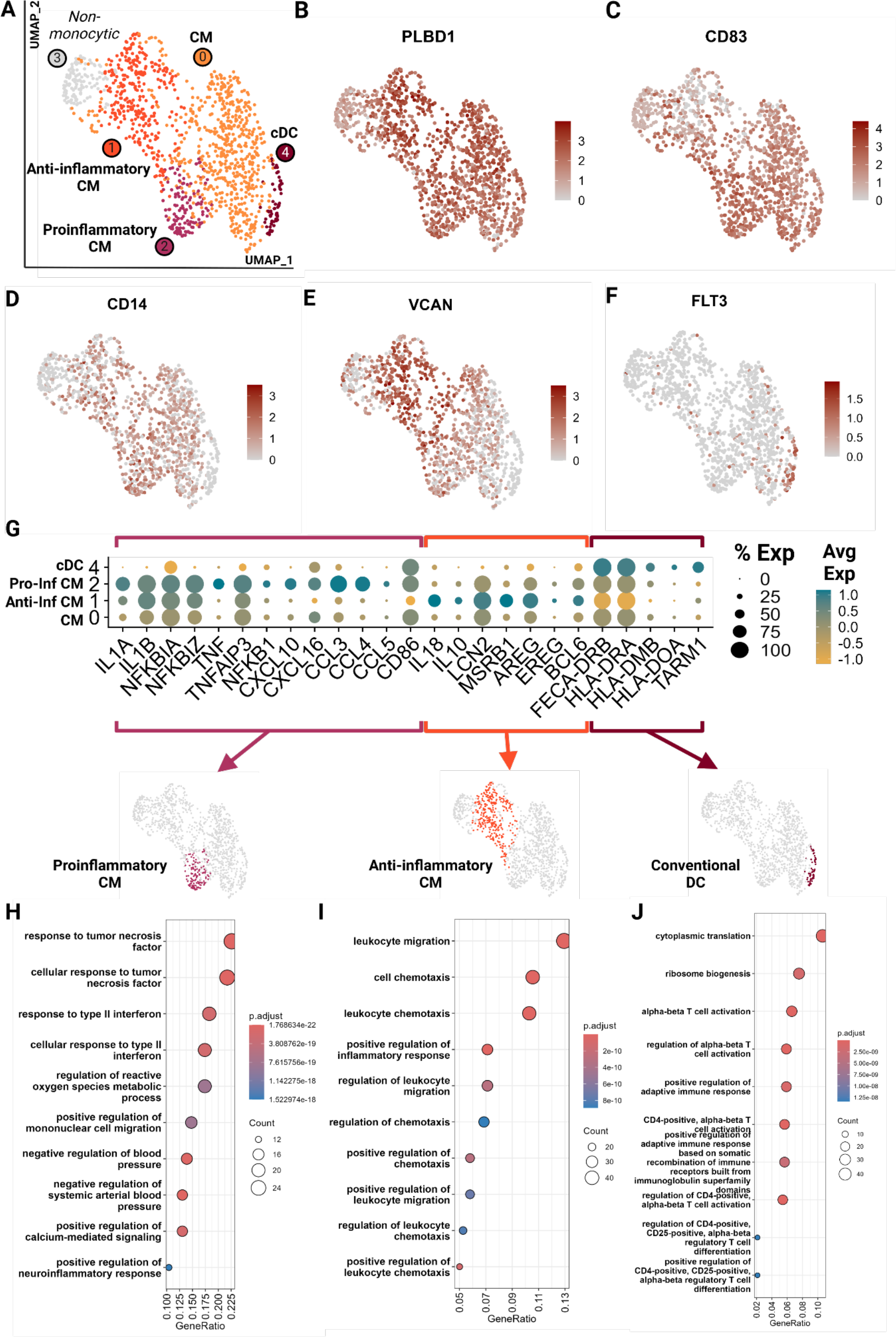
Feline circulating monocytes include classical monocytes with more differentiated clusters with traditional proinflammatory and unconventional anti-inflammatory phenotypes. Unsupervised clustering revealed 5 clusters. (A) UMAP of scRNA-seq atlas of monocytes. (B- F) UMAP of monocytes colored by expression levels of canonical markers. (G) Dot plot of genes of monocytic phenotypes across the 4 monocytic clusters. (H-J) Dot plots of top GO biological process terms called based on sets of positive differentially expressed genes identified via Seurat *FindMarker* function (Wilcoxon rank sum, *Adj P <0.05*). Classical monocyte, CM; Conventional dendritic cell, cDC; Pro-inf, proinflammatory; Anti-inf, anti- inflammatory.

In-depth assessment of differentially expressed genes across the four monocytic clusters identified signatures that further characterized these cell types. Cluster 2 demonstrated a proinflammatory phenotype characterized by upregulation of classic proinflammatory cytokines (*IL1A*, *IL1B*), *TNF/NF-KB* pathway (*TNF*, *TNFAIP3, NFKBIA*, *NFKBIZ*, *NFKB1*) and M1-macrophage-associated proinflammatory cytokines (*CXCL10, CXCL16, CCL3, CCL4 and CCL5*) **<Fig 7G,H, Supplementary File 7,8>** (20,50–54). Cluster 1 gene ontology revealed an upregulation of leukocyte migration and chemotaxis terms in addition to upregulation of proinflammatory cytokines *IL1A* and *IL1B* **<Fig 7G,I, Supplementary File 7,8>**. However, these cells also lack *TNF* expression and upregulate anti-inflammatory cytokines *IL10* and *IL18* and proteins which promote anti-inflammatory phenotypes in macrophages such as *LCN2* and *MSRB1* (55,56) **<Fig 7G>**. They have the lowest expression of *CD86*, a T cell co-stimulatory molecule of antigen presenting cells, which is associated with an anti-inflammatory phenotype (57). Most strikingly, these cells were found to upregulate genes similar to a novel monocyte cluster found in SARS-CoV-2 infected and recovered patients. These genes include *AREG*, *EREG*, *BCL6*, *IL10,* and *IL18* which have been described as anti-inflammatory tissue repair genes **<Fig 7G>** (58,59). Thus, these cells were designated as anti-inflammatory, although they do not follow the traditional M2 polarization phenotype described in human monocytes (60). Our data suggests a novel classical monocyte subtype with an anti-inflammatory phenotype.

Cluster 0 (CM) appears the least differentiated with low expression of inflammatory cytokines and the top GO terms include more cellular rather than immune related functions **<SUPPLEMENTARY File1 S8>**. Cluster 4 (cDC) showed an upregulation of and the GO terms associated with leukocyte activation and T cell activation via TCR contact with antigen bound MHC **<Fig 7G,J, Supplementary File 7,8>**. Given the strong antigen presentation phenotype, the annotation of cDC was further corroborated.

### 2.7 Feline circulating neutrophils

We identified 2 major neutrophil clusters **<Fig 8A>** based on the expression of canonical neutrophil markers including *CSF3R* and *ELANE* as well as mature neutrophil functional markers vimentin (*VIM)*, thioredoxin (*TXN)*, *S100A9* and *S100A12*, suggesting both clusters are comprised of mature neutrophils (25,61,62) **<Fig 8B-G>**. Cluster 1 exhibited several notable differences compared to cluster 0. Differential gene expression analysis accompanied by gene ontology showed an upregulation in cluster 1 of apoptotic signaling and activated neutrophil functions such as response to lipopolysaccharide and *IL-2* production **<Fig 8H, Supplementary File 9,10>**. Thus, we annotated this cluster as activated neutrophils. When investigating top differentially expressed genes, we noted a higher expression of *IL1A* and *TNFAIP3*, suggesting a proinflammatory phenotype (63) **<Fig 8I,J>**. Similarly, cluster 1 had an increased expression of *VCAN*, which is upregulated in the skin in response to damage from ultraviolet light B and reactive oxygen species (64). Lastly, cluster 1 exhibited lower RNA counts per cell, which is consistent with findings in murine neutrophils where the number of genes and counts of RNA decrease with neutrophil differentiation, maturation and eventual activation and function (65). Since a pathway driving this clustering could not be resolved, we examined an interferon (IFN) module score, which is a composite score based on expression levels of interferon stimulated genes **<Fig 8N>**. Cluster 1 did not show an increased IFN score but the adjacent cells in cluster 0 did **<Fig 8N>**. Interferon stimulation can delay neutrophil apoptosis, which could represent one of the potential driving factors for cells to move from cluster 0 to cluster 1 (66). This analysis is the first to resolve circulating neutrophil transcriptomic signatures in cats at the single cell level.

**Figure 8.**
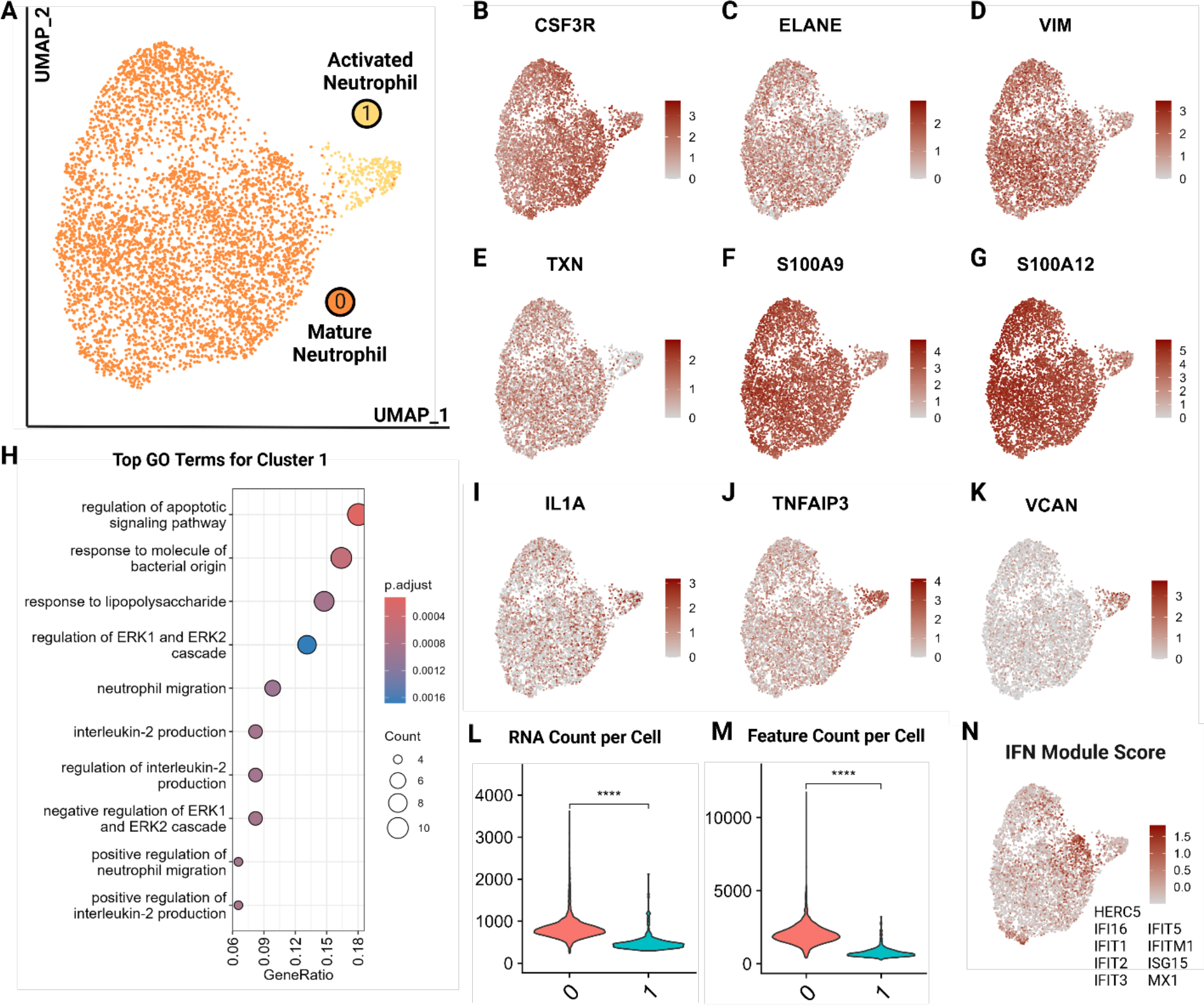
Feline circulating neutrophils separate into 2 clusters based on activation state. (A) UMAP of scRNA-seq atlas of neutrophils. (B-G) UMAP of neutrophils colored by expression levels of canonical and functional neutrophil markers. (H) Dot plots of top GO biological process terms in cluster 1 called based on sets of positive differentially expressed genes identified via Seurat *FindMarker* function (Wilcoxon rank sum, Adj *P <0.05*). (I-K) UMAP of neutrophils colored by expression levels of select differentially upregulated genes in cluster 1. (L,M) Violin plots of RNA counts and feature (unique RNA) counts per cell by cluster; (****) indicates P value less than 0.001 by T-test. (N) UMAP of neutrophils colored by interferon (IFN) gene composite score per cell; genes included are named in figure.

## 3 Discussion

The importance of the cat as a companion animal as well as a model for infectious diseases warrants an in-depth characterization and understanding of the feline immune system. Our study is one of the first to characterize feline immune cell subpopulations at the single cell level. We utilized 5’ single cell RNA-sequencing with V(D)J analysis to resolve the heterogeneity of CD5^+^ enriched peripheral blood immune cells. We resolved populations of T cells, neutrophils, B cells and monocytes. Additionally, this atlas is the first to annotate the single cell transcriptome of rarer cell types, plasmacytoid dendritic cells and mast cells, which are seldomly captured in single cell atlases (67,68).

Among T cells, we were able to resolve different populations of naive cells including a significant number of *CD8A*/*CD8B* naive cells. We further identified T effector cells and terminally differentiated cytotoxic effectors. The transcriptomic gradient of T cell differentiation was found to fit the paradigm seen with other species (23). Additionally, we captured feline γδ T cells for the first time. Upon investigation of effector cells, we were able to demonstrate known T helper effector phenotypes among the most differentiated effectors. Of note, the Treg phenotype was found to be the most prominent. The importance of Treg populations has been studied in cats, particularly in the context of feline mammary carcinoma (69–71). Our study corroborates the large presence of this phenotype and the potential importance of Treg cells in feline immunology.

We performed a cross species integrative analysis of T cells to take advantage of the growing database of single cell RNA data. We found relatively high conservation of T cell subtypes along an effector gradient with relatively equitable representation of veterinary species (horse, dog, pig) and humans with the cat. This analysis also revealed the potential negative impacts of CD5^+^ enrichment for T cells. Effector clusters with high *CD5* expression were found to be underrepresented in the cat. Thus, a broader T cell selection protocol would be necessary in future studies to further investigate effector cells in cats.

In addition to characterizing T cells based on their transcriptome, we analyzed TCR expression on a single cell level for the first time in the cat. Cytotoxic CD8^+^ T cells differed from other αβ T cell subsets in multiple aspects. In addition to the expected TRA and TRB transcripts, a high percentage of cytotoxic CD8^+^ T cells expressed a productive TRG transcript. Given that this was not observed in naïve CD8+ T cells, we attribute this phenomenon to differential transcriptional regulation of naive and effector T cells. Compared to other αβ T cell subsets, cytotoxic T cells also exhibited divergent TRAV and TRBV gene usage but not TRGV gene usage. Instead, the junctional length of TRG transcripts was skewed towards a highly conserved16 aa motif. This finding is of potential significance for clonality testing, an adjunct method for differentiating reactive from neoplastic lymphoid proliferations. If the majority of cytotoxic CD8^+^ T cells harbor a length restricted TRG rearrangement, then reactive CD8^+^ dominant T cell proliferations might exhibit reduced TRG diversity, which could result in a higher false positive rate. Of note, since other αβ T cell subsets did not express a significant number of productive TRG transcripts, it is unknown if similar length restricted rearrangements occur in these subsets. A previous study examining the feline TCR transcriptome across thymus, spleen and mesenteric lymph nodes did not identify any TRG length restrictions (13). However, the previous study aimed to identify global rearrangement patterns and hence utilized a bulk sequencing strategy, which is not suited to resolve repertoire characteristics of individual T cell subsets.

A cluster analysis of TCR sequences did not identify T cell subsets with invariant TRA or TRB chains. A previous study utilizing scRNA-seq with VDJ sequencing in dogs identified MAIT-like cells based on restricted TRAV gene usage and restricted junctional length (19). Although cytotoxic T cells in the present study exhibited differential TRAV gene and, to a lesser extent, differential TRBV gene usage, there was no obvious junctional length restriction or dominant amino acid motif. The fact that we were unable to identify MAIT-like cells in cats could reflect their true absence or represent an artifact of CD5-based enrichment, which might have resulted in the exclusion of CD5^-^ or CD5^low^ subsets. Additional studies with more animals, inclusion of different tissue types and without selection bias are needed to elucidate the existence of innate-like alpha/beta T cells in cats.

Among the myeloid cells, we resolved 3 clusters of classical monocytes and 1 cluster of conventional dendritic cells. Of the three clusters of monocytes, we found two to be differentiated in a somewhat polarized manner- pro-inflammatory versus anti-inflammatory. The anti-inflammatory population was unconventional in that it had an upregulation of classic pro-inflammatory cytokines *IL-1A* and IL*-1B* but also an upregulation of genes associated with a unique and novel monocytic cluster found in human COVID-19 patients such as *AREG, EREG, IL10*, *IL18* and *BCL6.* (59,72). Sub clustering of neutrophils identified a larger mature cluster and a smaller exhausted/activated cluster which showed upregulation of pro-inflammatory and apoptotic genes. Thus, analysis of these myeloid cells revealed the presence of conventional and non-conventional subtypes, providing a footing for feline circulating myeloid immunology.

Currently, T cell subtypes have only been described at the flow cytometric level, and with a focus on *CD4/CD8* populations due to paucity of species-specific reagents. Our data indicates the presence of unique subtypes of peripheral blood immune cells including αβγ CD8^+^ cytotoxic cells, *CD8A/CD8B* naive T cells, plasmacytoid DC, mast cells, conventional DCs, anti-inflammatory monocytes and mature and end stage neutrophils. Our study is the first of many necessary for the further characterization of the feline immune system. Taken together, it has increased our understanding of feline circulating immune cells, including T cell subtyping and capture of rare populations previously undescribed in the cat.

## 4 Materials and Methods

### 4.1 Samples and Sorting

Whole blood was obtained from four healthy, female, domestic shorthair cats (13-101/9 years, 18- 003/6 years, 21-005/1 year, 22-007/6 months) housed at the UC Davis Nutrition and Pet Care Center in accordance with the UC Davis Policy and Procedure Manual section 290-30 and Animal Use and Care protocol #22150. Blood was collected in purple top EDTA tubes and immediately processed in one batch without freezing until completion of the reverse transcription step during library preparation. Peripheral blood mononuclear cells (PBMC) were isolated from whole blood by density gravity centrifugation. Briefly, 5 mL of whole blood was layered on top of 10 mL Histopaque 1077 (density 1.077g/mL, Sigma-Aldrich, Burlington, MA, USA) in a 50 ml tube. The samples were centrifuged at 850 revolutions per minute (RCF) for 30 minutes at room temperature (RT) without break. After centrifugation, the PBMC layer at the plasma-histopaque interface was aspirated and transferred to a clean 15 ml tube. The PBMCs were then washed twice with 14 ml PBMC isolation media (500 mL Hanks’ Balanced Salt Solution (HBSS), 15 mL Fetal Bovine Serum (FBS), 2 mL EDTA) by centrifuging at 850 RCF for 10 minutes at RT. Following the second washing step, the PBMC pellet was resuspended in 200 μl modified flow buffer solution (MFB, 500 ml phosphate buffer saline (PBS), 0.467g EDTA tetrasodium salt and 1% Horse serum).

The isolated PBMCs were first incubated with a primary anti-feline CD5 monoclonal antibody (clone FE 1.1.B11-IgG1-Azide, Peter F. Moore Leukocyte Antigen Biology Laboratory, UC Davis, CA, USA) or a non-cross reacting canine antibody (negative control) for 30 min at RT. This was followed by a wash step (addition off 3 mL of MFB, followed by centrifugation at 400 RCF for 3 min at RT and resuspension in 100 - 200 μl of supernatant), and incubation with a secondary horse-anti-mouse FITC antibody (1:99 in MFB) for 15 minutes at RT in darkness. Finally, the stained PBMC was washed again (as above), resuspended in 500 μl of MFB and stained with DAPI (1 μg/mL). Cells were sorted at the UC Davis Comprehensive Cancer Center Flow Cytometry Core. Cells were initially gated based on FSC vs. SSC to exclude debris and aggregates. Singlet gating was then applied based on SSC to exclude doublets, followed by discrimination of live and dead cells based on DAPI staining and subsequent gating for CD5 positive (FITC positive) cells <SUPPLEMENTARY File1 S11>.

### 4.2 Single-cell 5’ RNA and V(D)J library preparation, sequencing and mapping

Libraries were constructed using the Chromium Next GEM Single Cell 5’ Kit v2 (10x Genomics). All four feline T cell receptor loci were amplified using the Chromium Single Cell V(D)J Enrichment Kit according to the manufacturer’s instructions but with custom reverse primers specific for the feline orthologs **<SUPPLEMENTARY File1 S12>**. Separate libraries were prepared for the transcriptome, alpha/beta TCRs, and gamma/delta TCRs **<SUPPLEMENTARY File1 S12>**. Libraries were sequenced using the Illumina NovaSeq S4 platform using 150 paired-end reads to a target read depth of 30,000 reads per cell. Raw FASTQ sequencing data were processed by the UC Davis Bioinformatics Core using 10X CellRanger version 7.1 for alignment to the cat reference genome (*Felis_catus_9.0*). To improve the quality of annotation of the *Felis catus* genome, we converted unmapped ENSEMBL IDs to human homologs from BioMart for greater interpretative power. Gene mapping was performed in the following order: (1) Mapping of one-to-one orthologs from cat to human (2) Mapping of one-to-many and many-to-many was performed by choosing a representative cat gene based on the following parameters, in respective order [orthology_confidence, gene_order_confidence, Whole_Genome_alignment, %_query_identical_target, %target_identical_query]. Additionally, specific unmapped genes were manually annotated or filtered **<SUPPLEMENTARY File1 S13>**. Further analysis was then continued using Seurat v5 (73).

### 4.3 Count matrix pre-processing and quality control

Filtered feature matrices for the 4 samples were independently normalized and scaled using the function *SCTransform* v2 (glmGamPoi) (74). Dimensionality reduction (determined using knee identification in elbow plot), k-means clustering (resolution determined using clustree package) and neighbor identification (74–76) was performed. Low quality clusters were identified and filtered based on low gene counts, high mitochondrial read percentage and lack of biologically relevant markers. To further ensure quality, doublet identification was performed using package *DoubletFinder* according to package vignette and ambient RNA decontamination was performed using the *DecontX* package as per package tutorials adapted to the data (77,78).

### 4.4 Integration and Downstream analysis

Objects were integrated by sample using the Seurat native reciprocal principal component analysis (RPCA) (79). The embeddings were utilized for Uniform Manifold Approximation and Projection (UMAP) and subsequent clustering and neighbor identification as described prior. Differential gene expression analysis was performed using each cluster identified by the Seurat function *FindAllMarkers* (Non-parametric Wilcoxon Rank Sum with adjusted P value <0.05). Gene set enrichment analyses including over-enrichment analysis (ORA) of gene ontology (biological process) was performed using *ClusterProfiler* on human homologized genes (adjusted P value <0.05) (80). Additional visualizations and associated statistics were performed using ggplot2 and gg supplementary packages (81). Figures were illustrated on Bio Render.

### 4.5 Cell type sub clustering

Cell type sub-clustering (example: T cell sub-clustering) analysis was performed in the same manner as described prior. Pseudotime values were calculated using the UMAP embeddings produced via Seurat with RPCA integration using package *monocle3* (82). Gene module scores were calculated via the Seurat function *AddModuleScore*.

### 4.6 Cross species T cell integrative analysis

Published single cell RNA-sequencing datasets were obtained from NCBI GEO Accession for 4 species including dog (*Canis familiaris*: GSE225599 (18)), human (*Homo sapiens*: https://support.10xgenomics.com/single-cell-gene-expression/datasets/3.0.0/pbmc_10k_v3), horse (*Equus caballus*: GSE148416 (22)) and pig (*Sus scrofa*: https://data.nal.usda.gov/dataset/data-reference-transcriptomics-porcine-peripheral-immune-cells-created-through-bulk-and-single-cell-rna-sequencing. (21)). All species genes were mapped to human homologs as described prior for the cat. Only homologized shared genes between all 5 species were utilized for analysis. T cell clusters annotated by authors were validated using canonical T cell markers (*CD3E*, *CD3D*, *TCF7*, *LCK*, *ITK*). T cells were integrated across species using Seurat canonical correlation analysis (CCA) and downstream analysis proceeded in the same manner as described prior for global object analysis.

### 4.7 VDJ analysis

Contigs were mapped to available IMGT reference genes (as of February 2023) using Cell Ranger version 7.1. All analyses were based on the ‘all_contig_annotations.csv’ file using the R packages tidyverse, circlize and ggseqlogo (83–85). Cluster analysis was performed as previously described (86).

### 4.8 Data availability

All raw and processed data have been submitted to NCBI GEO and are available under GSE267355.

## Acknowledgements

This project was funded by grant #2022-24-F from the Center for Companion Animal Health, University of California, Davis. The project was supported by the University of California Davis Flow Cytometry Shared Resource Laboratory with funding from the NCI P30 CA093373 (Comprehensive Cancer Center) and S10 OD018223, with technical assistance from Bridget McLaughlin, Jonathan Van Dyke and Ashley Karajeh. The sequencing was carried out by the DNA Technologies and Expression Analysis Core at the UC Davis Genome Center, supported by NIH Shared Instrumentation Grant 1S10OD010786-01. We thank Jie (Jessie) Li from the UC Davis Bioinformatics Core for data analysis and Kristy Harmon from the Leukocyte Biology Antigen Laboratory for her flow cytometry support.

