## Supplementary File 1 for "Single cell RNA-sequencing of feline peripheral immune cells with V(D)J repertoire and cross species analysis of T lymphocytes"

### Slide 1
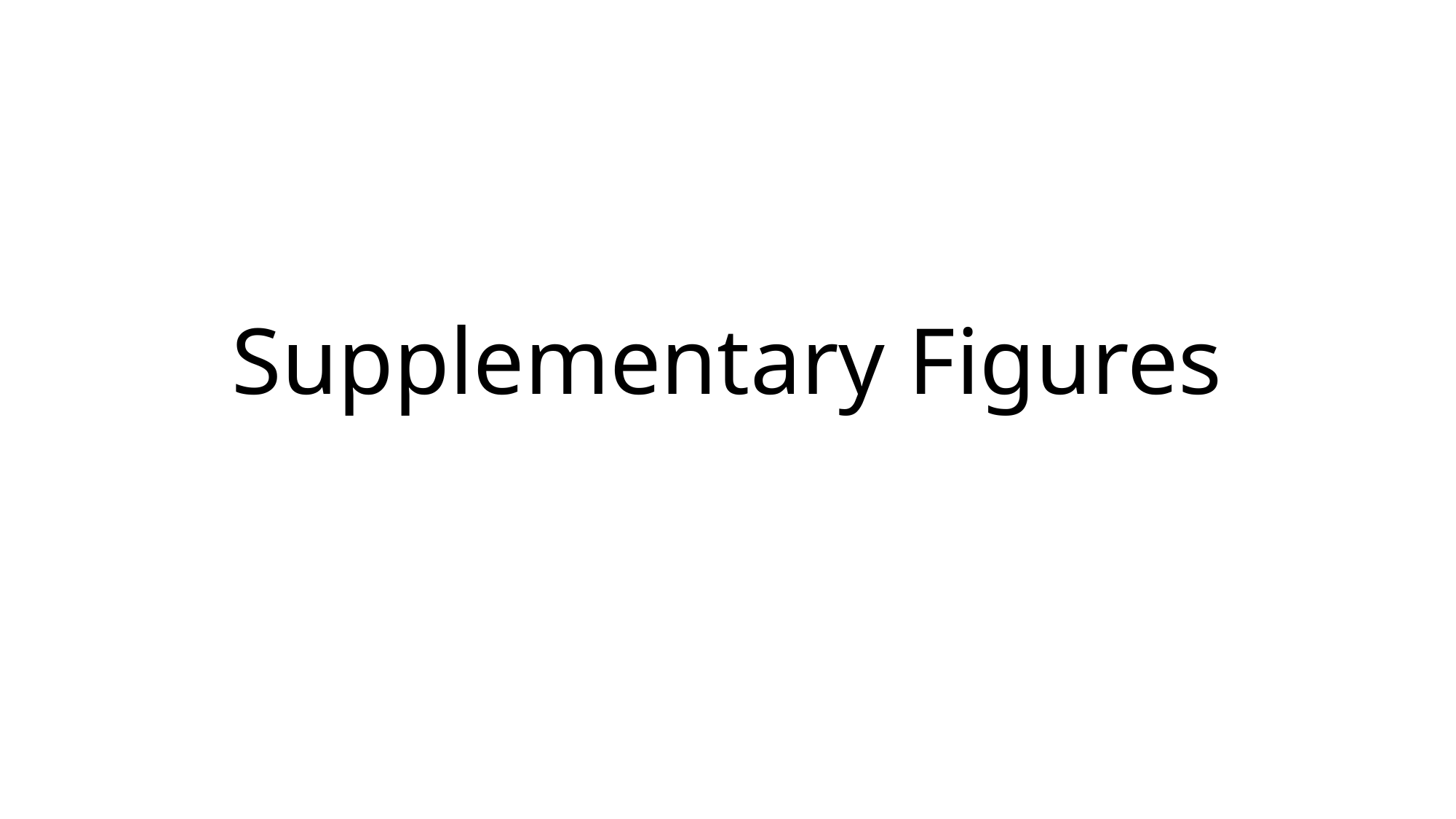

#### Supplementary Figures

### Slide 2
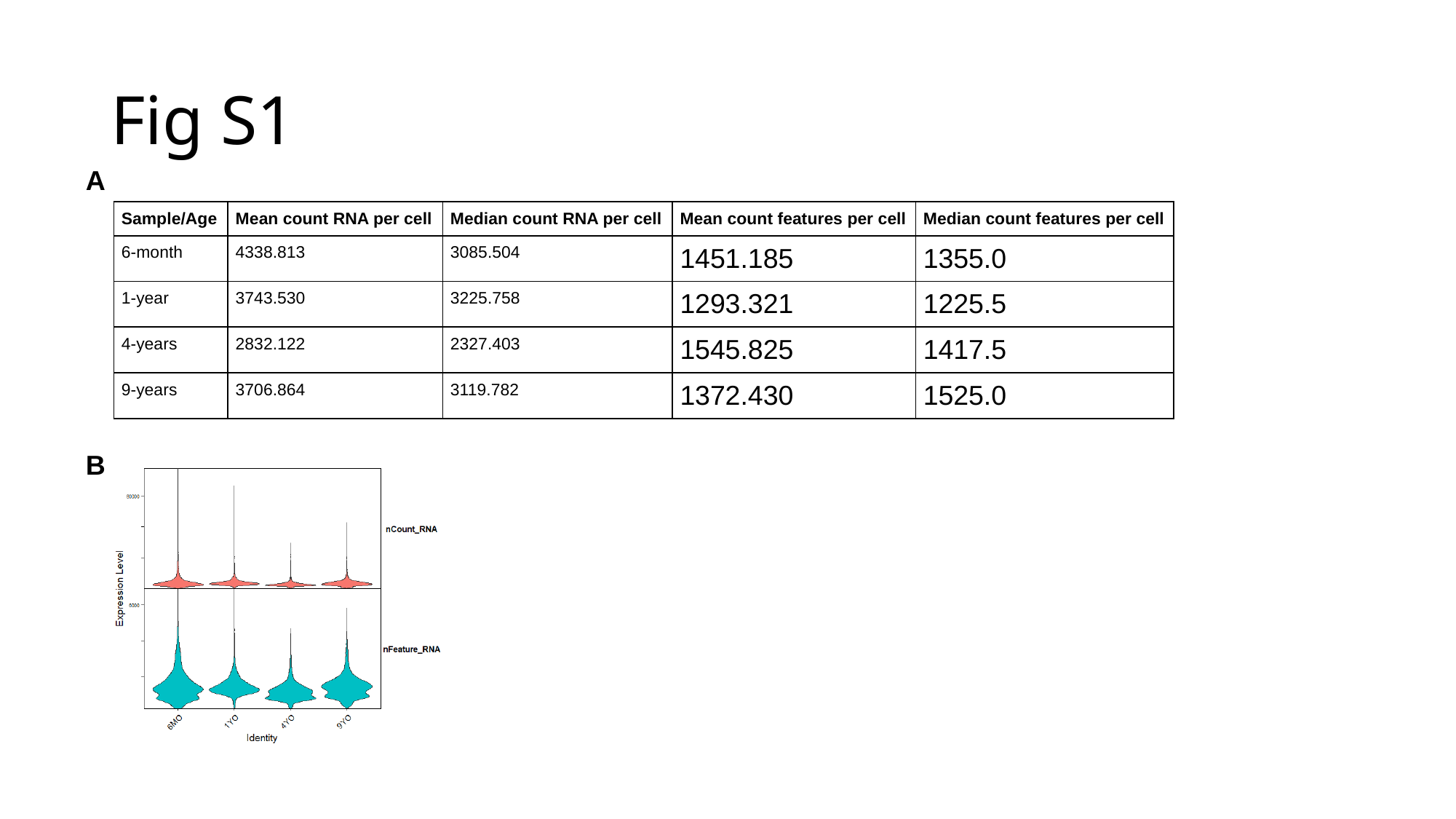

#### Fig S1
A
| Sample/Age | Mean count RNA per cell | Median count RNA per cell | Mean count features per cell | Median count features per cell |
| --- | --- | --- | --- | --- |
| 6-month | 4338.813 | 3085.504 | 1451.185 | 1355.0 |
| 1-year | 3743.530 | 3225.758 | 1293.321 | 1225.5 |
| 4-years | 2832.122 | 2327.403 | 1545.825 | 1417.5 |
| 9-years | 3706.864 | 3119.782 | 1372.430 | 1525.0 |
B

### Slide 3
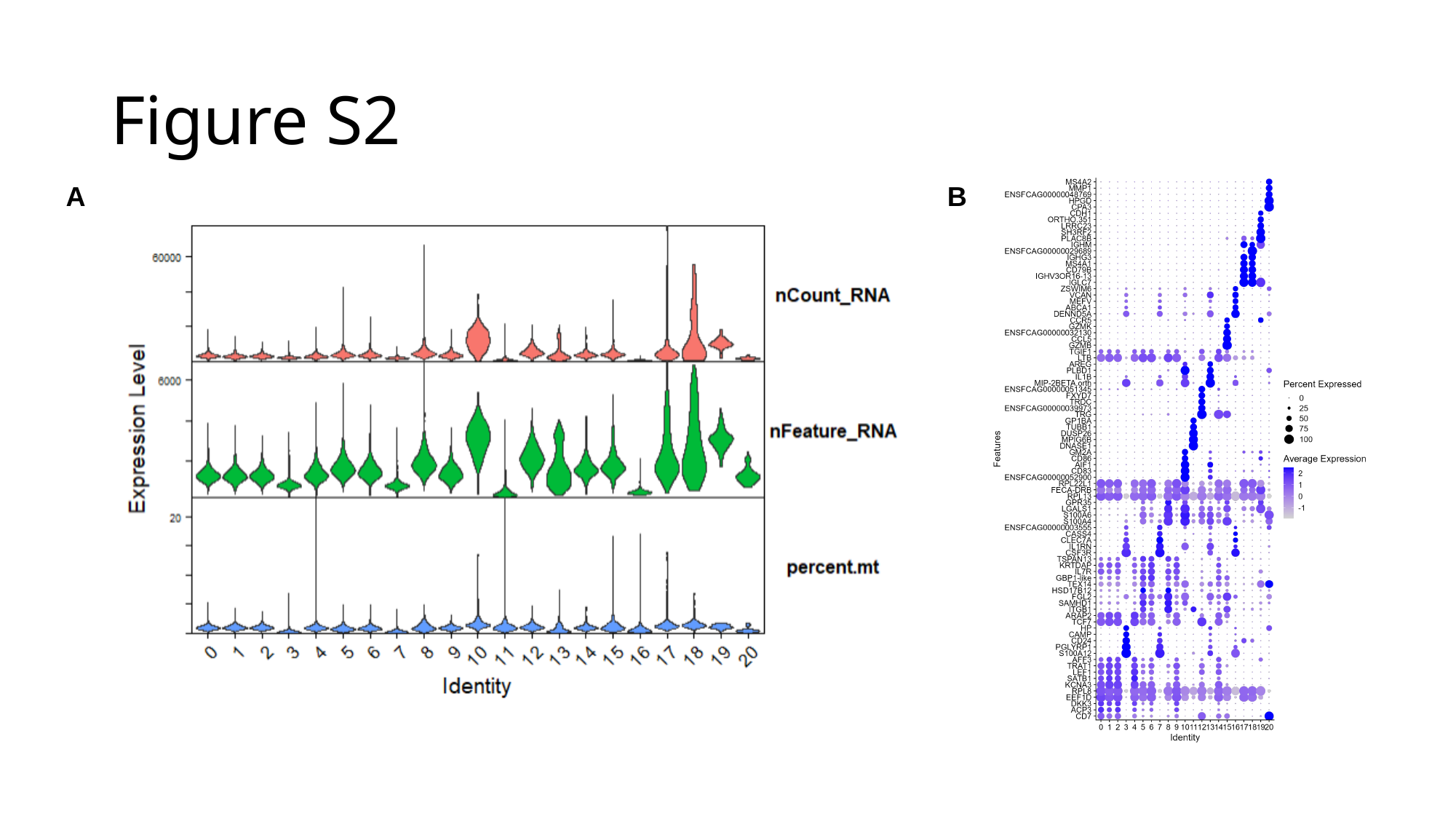

#### Figure S2
A
B

### Slide 4
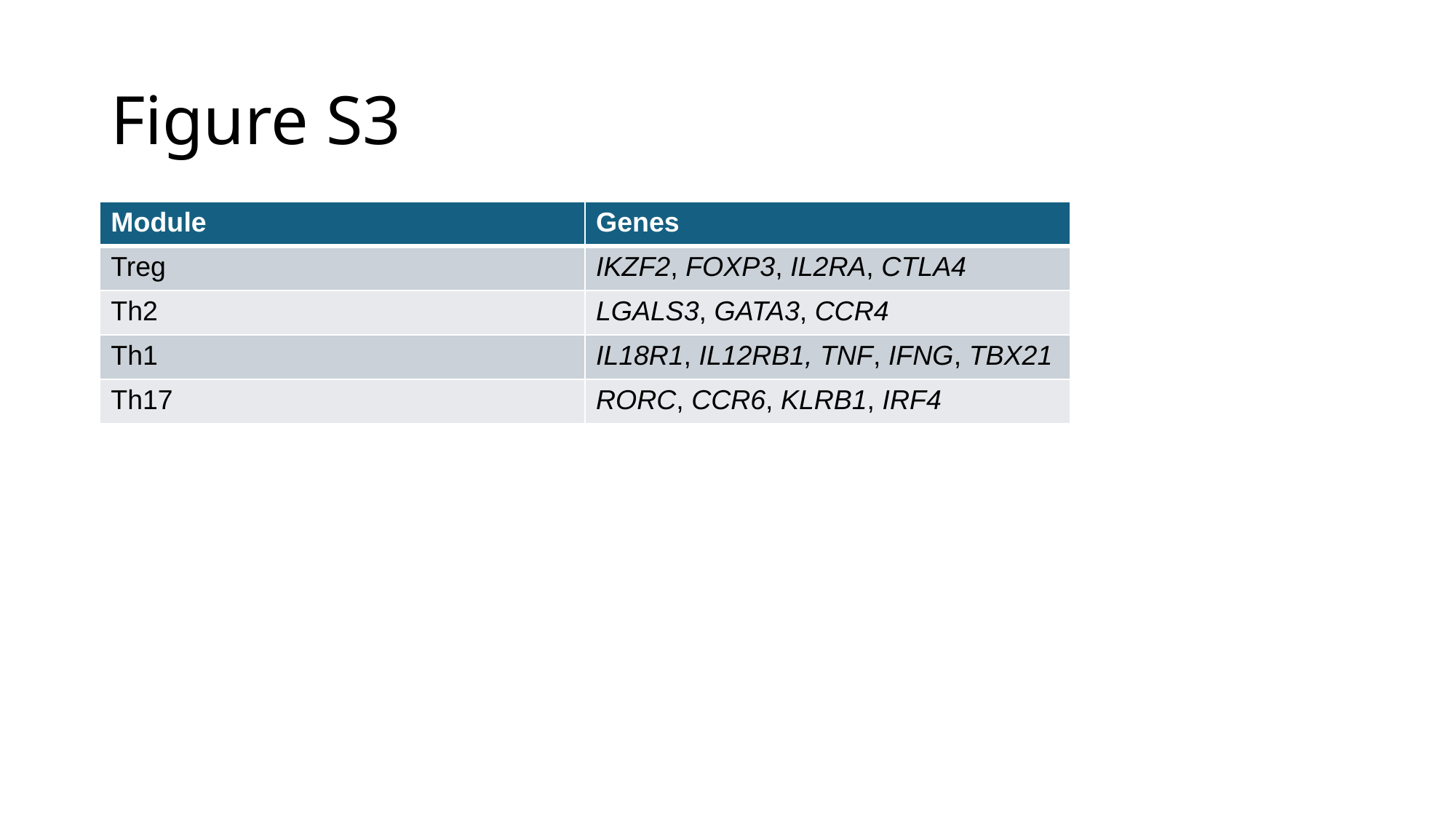

#### Figure S3
| Module | Genes |
| --- | --- |
| Treg | IKZF2, FOXP3, IL2RA, CTLA4 |
| Th2 | LGALS3, GATA3, CCR4 |
| Th1 | IL18R1, IL12RB1, TNF, IFNG, TBX21 |
| Th17 | RORC, CCR6, KLRB1, IRF4 |

### Slide 5
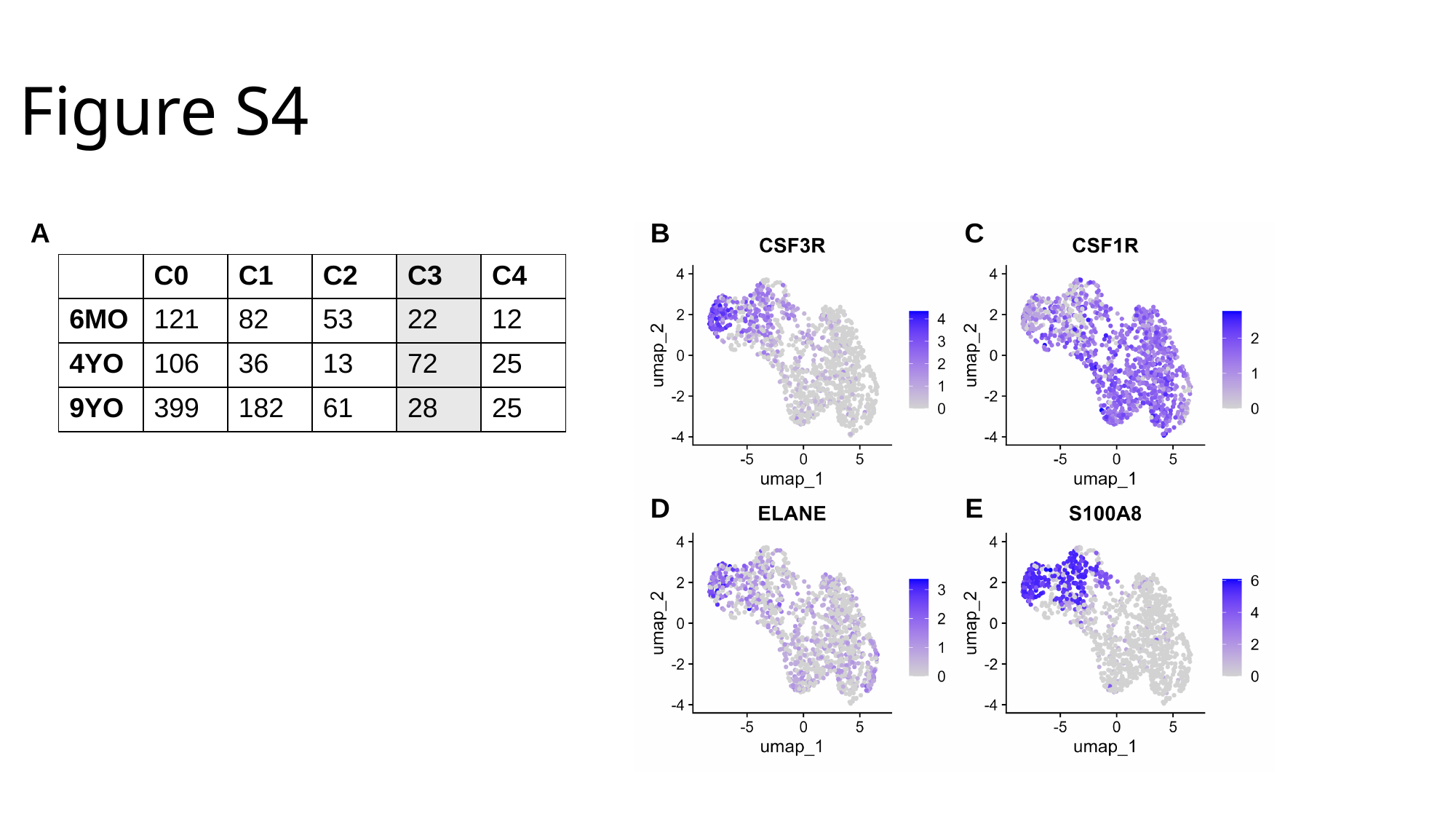

#### Figure S4
A
C
B
| | C0 | C1 | C2 | C3 | C4 |
| --- | --- | --- | --- | --- | --- |
| 6MO | 121 | 82 | 53 | 22 | 12 |
| 4YO | 106 | 36 | 13 | 72 | 25 |
| 9YO | 399 | 182 | 61 | 28 | 25 |
D
E
