## Supplementary figures and images for "Single cell RNA-sequencing of feline peripheral immune cells with V(D)J repertoire and cross species analysis of T lymphocytes"

### Supplementary File 11

#### Slide 1
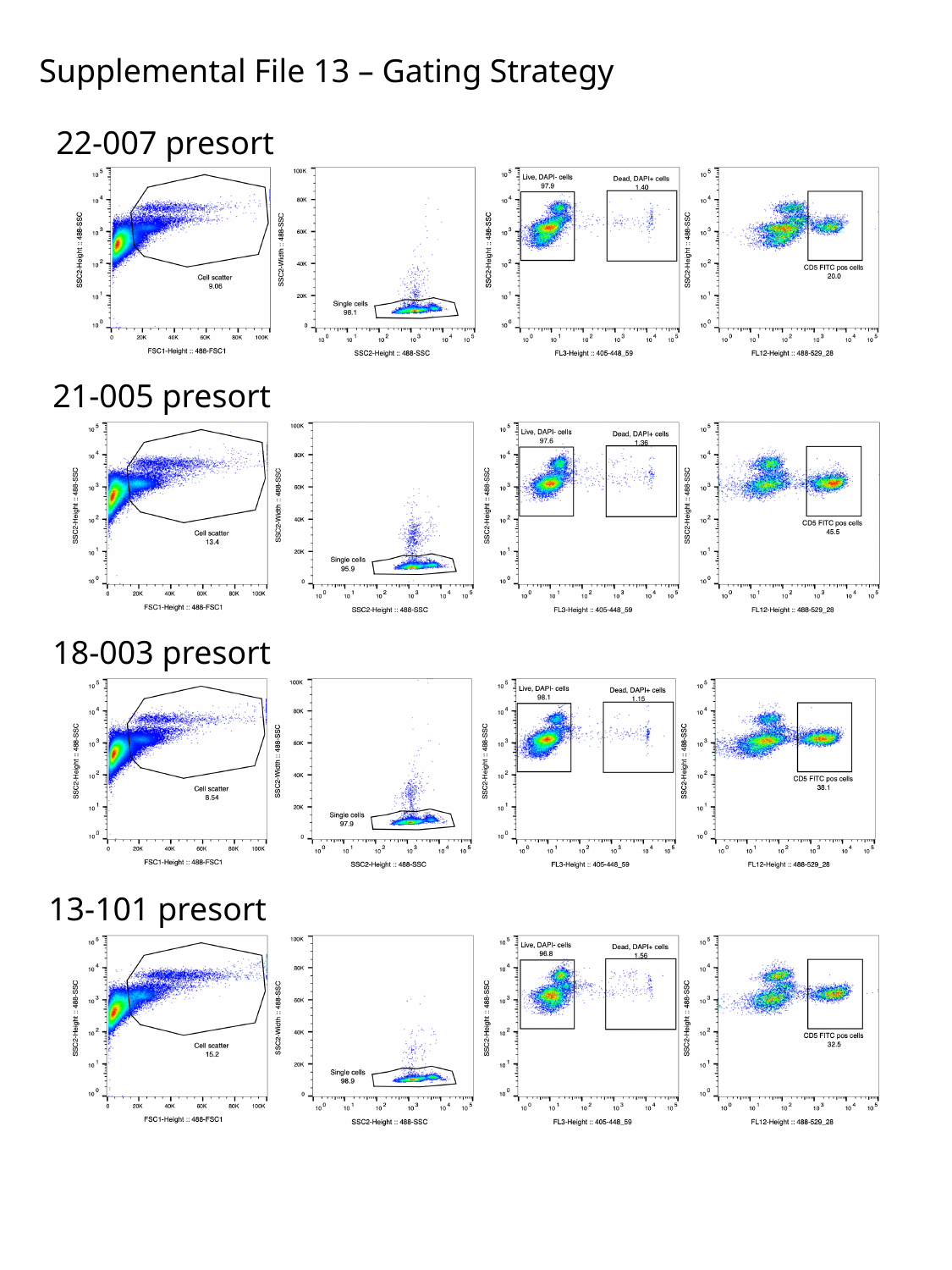

Supplemental File 13 – Gating Strategy
22-007 presort
21-005 presort
18-003 presort
13-101 presort
